# Gene network inference and master regulator analysis identifies the estrogen-related receptor gamma (ERRγ) as a therapeutic target for alcohol use disorder (AUD)

**DOI:** 10.1101/2025.11.15.688629

**Authors:** Irene Lorrai, Riccardo Maccioni, Itzamar Torres, Roberta Puliga, Jorge Marquez-Gaytan, Federico M. Giorgi, Vez Repunte-Canonigo, Pietro Paolo Sanna

## Abstract

Differential gene expression is often inadequate to predict the activity of transcription factors and their contribution to the phenotypes associated with specific gene expression states. Here we used a systems biology approach based on gene network inference and master regulator analysis (MRA) to identify candidate drivers of the gene network dysregulations in the prefrontal cortex (PFC) of human subjects with a history of alcohol dependence. The estrogen-related receptor gamma (ERRγ) gene ESRRG, an orphan nuclear receptor protein that acts as a transcription activator, emerged as a high-ranking Master Regulator (MR) based on the expression of its targets and was selected for functional validation due to its translational and druggability potential. The ERRγ agonist, GSK4716, reduced alcohol drinking in the mouse binge drinking paradigm of drinking in the dark (DID) and in both non-dependent mice as well as in mice made dependent by chronic intermittent vapor exposure (CIE). GSK4716 also prevented alcohol-conditioned place preference without affecting saccharin intake or mouse locomotion. Similarly, in rats, GSK4716 reduced operant oral alcohol self-administration in non-dependent and dependent (by CIE) rats under fixed and progressive ratio schedules of reinforcement. Overall, these results support the efficacy of transcriptome-wide gene regulatory network approaches for the identification of key druggable regulators of long-term transcriptional adaptations that sustain the molecular and behavioral pathology of alcohol dependence and identify ERRγ as a regulator of excessive alcohol drinking and seeking, and a therapeutic target for AUD.

## Introduction

We used a systems biology strategy for the reconstruction and interrogation of genome-wide gene regulatory networks to identify drivers of the gene network dysregulations associated with a history of alcohol dependence in the pre-frontal cortex (PFC) in human subjects [1]. Master regulator analysis (MRA) methods aim to identify transcription factors (TFs) and their modulators that drive the transition between two phenotypes and sustain the resulting phenotype [2–4]. Differential gene expression is frequently inadequate in predicting the TF’s activity and its contribution to the implementation of the gene expression signatures associated with a specific disease condition that drive the phenotype [2–4] such as excessive drinking [5]. To address this problem, MRA algorithms infer TF activity from the transcriptional dysregulation of its target genes or regulon [2–4]. Here we applied this approach to the identification of genes driving excessive alcohol drinking in humans with a history of alcohol dependence.

We used the *corto* algorithm and pipeline [4] to identify key genes, master regulator genes (MRs), contributing to the PFC gene expression signature associated with a history of alcohol dependence in a RNA-Seq gene expression dataset [1]. Interrogation of this dataset identified an unbiased set of candidate MRs whose activity differentiates between the conditions under study – alcohol-dependent/control – and thus are likely to be causally responsible for regulating the PFC transcriptional signatures associated with alcohol dependence. Among them, we selected the estrogen-related receptor gamma (ERRγ) for functional validation on the basis of prioritization criteria, including its rank in the inference list and its translational and druggability potential.

ERRγ, together with ERRα and ERR*β*, belongs to the estrogen-related receptor (ERR) subfamily of the group 3 steroid nuclear receptor superfamily (NR3B1-3). These receptors share significant sequence homology with estrogen receptors (ERs); however, estrogens are not their endogenous ligands [6]. ERRs show constitutive activity and function as transcriptional regulators even in the absence of ligand binding [7]. In addition, several transcriptional co-activators have been identified; among these the peroxisome proliferator-activated receptor gamma co-activators 1-alpha (PGC1-α) and PGC1-*β* [8] which enhance ERRs’ transcriptional activity of genes implicated in mitochondria biogenesis and, more broadly, cellular energy metabolism [7, 9].

ERRγ is localized in the nucleus of neurons in the central nervous system (CNS) [10, 11] and it is widely distributed in the adult brain [12–14]. A growing body of literature reveals an important role for ERRγ in various physiological and pathological states [15–21], including neurodegenerative disorders such as Alzheimer’s disease (AD) [22] and Parkinson’s disease (PD) [23]. Although ERRγ is an orphan receptor, in the past few years, several small molecules have been synthetized to better define the pharmacological and functional profile of ERRγ [24, 25].

Here we show that acute administration of the ERRγ agonist, GSK4716, significantly reduces alcohol consumption in rodents across multiple validated behavioral paradigms of alcohol use disorder (AUD), indicating that ERRγ is a key regulator of excessive alcohol drinking and seeking, and represents a promising therapeutic target for AUD.

## Materials and Methods

### Gene network analyses

We performed master regulator analysis (MRA) using the *corto* algorithm and pipeline [4], running the *corto* package version 1.2.4 on R version 4.5.1. As a transcriptome-wide quantitative gene expression dataset we used the Microarray (Illumina HumanHT-12 v 3.0) human brain collection deposited on Gene Expression Omnibus entry GSE29555 [1]. The signature was defined by contrasting 68 individuals with histories of alcohol dependence based on DSM4 criteria vs. 60 control samples extracted from post-mortem prefrontal cortexes (PFC) of human autopsy brain samples from the New South Wales Tissue Resource Centre at the University of Sydney [1]. We derived a human-based PFC gene regulatory network from the human GTEX [26] frontal cortex collection of 129 human samples using *corto* with default parameters and, as candidate master regulators (MRs), a curated list of human transcription factors (TFs) derived from Gene Ontology category DNA-binding transcription factor activity (GO:0003700). The MRA analysis used the aforementioned signature and regulon and was also executed with *corto* using default parameters and minimum regulon size of 15 (i.e., including only TFs with at least 15 targets with detected expression in the GTEX dataset).

In order to define the ESRRG (ERRγ) gene regulatory network we performed a correlation analysis between genetical and transcriptomical events using a modified DIGGIT pipeline [27]. Briefly, we collected 23 TCGA cancer datasets including gene-centered somatic mutation events to provide transcriptional perturbations to the gene regulatory network. The datasets included these tumor codes and cover the majority of public cancer data currently available: BLCA, BRCA, CESC, COAD, ESCA, GBM, HNSC, KIRC, KIRP, LGG, LIHC, LUAD, LUSC, OV, PAAD, PCPG, PRAD, SARC, SKCM, STAD, TGCT, THCA, THYM. Only somatic mutations affecting protein sequence were considered. Correlation between somatic events and the activity of the ESRRG regulon was calculated using DIGGIT [27]. The effect of the somatic mutation-carrying gene on ESRRG was calculated by integrating NESs across the pan-cancer TCGA dataset using the Stouffer method [28]. The layout of the DIGGIT graph is calculated using the Fruchterman-Reingold algorithm, a force-directed graph drawing method [29].

### Animals

Male C57BL/6J mice (6 weeks old, The Jackson Laboratories, USA) were either single or group housed according to the experimental design. Male Wistar rats (4 weeks old, Charles River, USA) were housed in pairs. All animals were kept in standard plastic cages under controlled temperature (21 ±1°C) and humidity (50±5%). Food and water were available ad libitum, except when specified otherwise. Behavioral experiments were conducted during the dark phase of the light/dark cycle. All procedures adhered to the National Institutes of Health guidelines for the “Care and Use of Laboratory Animals” and were approved by the Institutional Animal Care and Use Committee of The Scripps Research Institute.

### Drugs

The ERRγ receptor agonist GSK4716 (0, 1.25, 2.5, and 5 mg/kg) (Tocris, Minneapolis, MN, USA) was suspended in saline with 1% (w/v) Tween 80 and administered intraperitoneally (IP) 30 min prior to the behavioral experiments. The administration volumes were 10 ml/kg for mice and 2 ml/kg for rats.

## Experimental procedures

### Mouse experiments

#### Drinking the Dark paradigm

The acute effect of the GSK4716 on binge-like alcohol drinking was assessed in male C57BL/6J mice using the Drinking in the Dark (DID) paradigm conducted as described by Rhodes and colleagues [30, 31]. Briefly, three hours after lights-off, the water bottle of single-housed mice was replaced with a bottle containing 20% (v/v) alcohol and left in place for 2 hours. This procedure was repeated over three consecutive days during which mice acquired and stabilized their drinking behavior. Lastly, on the fourth day, C57BL/6J mice received an IP injection of either GSK4716 or vehicle before the start of the drinking session, which had a duration of 4 h. Alcohol intake was measured after the first and second 2 h interval. The amount of alcohol consumed was determined by weighing the bottles (± 0.01 g accuracy) immediately before and after each drinking session and expressed as grams of pure alcohol per kilogram of body weight (g/kg).

#### Non-drug reinforces in the Drinking in the Dark paradigm

To evaluate the effect of GSK4716 on non-drug reinforces, a separate cohort of male C57BL/6J mice were given access to saccharin [0.002% (w/v)] solution during 2 h sessions for three consecutive days. On Day 4, mice received an IP injection of GSK4716, or vehicle followed by a 4 h session with access to saccharin solution. Saccharin intake (ml/30g) was measured at two time points: after the first 2 h and at the end of the 4 h session, mirroring the procedure used in the alcohol experiment. The amount of solution consumed was determined by weighing the bottles (± 0.01 g accuracy) immediately before and after each drinking session.

#### Two-bottle choice paradigm and chronic intermittent exposure (CIE) to alcohol vapors

Male and female C57BL/6J mice were singly housed and exposed to the conventional 2-bottle alcohol [15% (v/v)] vs water choice (2BC) paradigm for 2 h over 5 consecutive days. Alcohol and water consumption were recorded by weighing the bottles before and after the drinking session. Subsequently, mice were divided into two balanced groups based on their alcohol and water intake during the baseline period. One group (dependent) was exposed to intermittent ethanol vapor (CIE, 16h ON, 9h OFF), and the other (non-dependent) to control air in identical chambers. Before being exposed to alcohol vapors, mice allocated in the dependent group were injected IP with a solution of alcohol (1.5 g/kg, 15% w/v in saline) containing 68.1 mg/kg pyrazole and immediately placed into alcohol vapor chambers (La Jolla Ethanol Research, CA, USA). Tail blood sampling for blood ethanol levels (BALs) determination was performed daily and targeted the 150–200 mg% range. Seventy-two hours following removal from the chambers, mice received access to water vs 15% (v/v) alcohol for 2 hours, and again over the next 4 days. The following week, mice were re-exposed to the alcohol vapor/control conditions and again tested for 2BC drinking for 5 days. Six total alcohol vapor cycles followed by 2BC were carried out. Mice were weighed every 4–6 days throughout the 2BC sessions and daily during the vapor exposure bouts.

#### Conditioned place preference in C57BL/6J mice

The ability of GSK4716 (5 mg/kg) to prevent the reinforcing effects of alcohol was assessed in male C57BL/6J mice using the conditioned place preference (CPP) test. The apparatus consisted of two compartments separated by a guillotine door, each with different visual and tactile cues to allow easily discrimination. The experimental protocol included three phases, as described elsewhere [32]. Briefly, on Day 1 (pre-conditioning phase), mice were allowed to freely explore the apparatus for 15 min to determine a baseline compartment preference. On Days 2-6 (conditioning phase), mice received an IP injection of either vehicle or GSK4716, and thirty minutes later an IP injection of either vehicle or alcohol [20% (v/v), 2 g/kg] and were confined to one compartment for 30 min. Eight hours later, mice received an alternate treatment and were confined to the opposite compartment. Treatment order was alternated across days to avoid sequence bias. On Day 6 (post-conditioning, test), mice were allowed to freely explore the entire apparatus for 15 min. An increase in time spent in the alcohol-paired compartment compared with pre-conditioning indicated conditioned place preference.

#### Spontaneous locomotor activity

Acute effect of GSK4716, at the same time and doses used in the DID experiments, was assessed on spontaneous locomotor activity in male alcohol-naive C57BL/6J mice. Briefly, three hours into the dark phase of the light/dark cycle, mice were injected with either GSK4716 or vehicle and placed into an empty box provided with infrared photo beams (Med Associates, St Albans, VT, USA). Locomotor activity was measured over a 2 h session.

### Rat experiments

#### Alcohol self-administration

##### Apparatus

Self-administration sessions were conducted in operant chambers enclosed in sound-attenuating and ventilated cubicles (Med Associates, St. Albans, VT, USA). The front panel of each chamber was provided with (i) two retractable and (ii) one liquid receptacle located between the two levers, and (iii) two stimulus lights and (iv) a sonalert mounted above each lever. The liquid receptacle was connected by polyethylene tubes to a syringe pump located outside the chamber. Responses on the right (active) lever determined the activation of the syringe pump and the delivery of 0.1 ml of alcohol 10% (v/v) into the receptacle. Alcohol delivery was paired to the illumination of a white light and turning on of a sound over 3 sec. Responses on the left (inactive) lever were recorded but did not have any scheduled consequences. *Two-bottle choice and Operant Training*. One week after acclimatization, male Wistar rats were exposed to the 2-bottle “alcohol (10%, v/v) vs water” choice paradigm with unlimited access (24 h) for 7 consecutive days. This procedure aimed to familiarize the rats with the taste of alcohol and promote its consumption. Subsequently, the alcohol bottle was removed, water was the only available fluid, and the rats were left undisturbed for 2 days. On the next day, the rats were trained to lever-respond for alcohol in operant chambers. On Day 1, the rats were exposed to a fixed ratio (FR) 1 (FR1) schedule of reinforcement for 10% alcohol (v/v) over a 12 h session. Water and food were provided in the chamber. On the following day (Day 2) the rats were left undisturbed in their home-cage. Alcohol self-administration resumed on Day 3, and session length reduced to 60 min, and subsequently 30 min over the next 2 days. On Day 5, a second inactive lever was introduced. Responses on this lever were recorded but had no programmed consequences. From this day onward, both the active and inactive levers were available, and the session length was set to 30 minutes for the remainder of the experiment, except as otherwise reported. *Fixed ratio 1 (FR1) schedule of reinforcement and chronic intermittent alcohol vapor exposure*. After 10 consecutive sessions of alcohol self-administration, rats were divided in two groups based on the number of lever-responses on the active lever and amount of self-administered alcohol over the last 3 days and then exposed to either air (non-dependent group) or alcohol vapors (dependent group). Alcohol dependence was achieved by exposing the rats to chronic, intermittent alcohol vapors (14 h ON, 10 h OFF, target BALs ~ 150-250 mg/dL, see [33]). After 4 weeks of vapor exposure, alcohol self-administration was resumed. The rats underwent 30 min alcohol self-administration sessions three-times a week (Monday, Wednesday and Friday) during acute withdrawal (6-8 h after vapors OFF). Non-dependent rats were tested on the same schedule. Following stabilization of lever responding, testing with GSK4716 on a 30-min FR1 self-administration session was conducted in both non-dependent and dependent rats. *Progressive ratio (PR) schedule of reinforcement*. After a few sessions of regular self-administration, non-dependent and dependent rats were tested under the progressive ratio (PR) schedule of reinforcement. Under these conditions, the response requirement necessary to receive a drop of alcohol increased progressively as follows: 1, 2, 4, 6, 9, 12, 15, 20, 25, 32, 40, 50, 62, 77, 95, 118, 145, 178, 219, 268, etc. according to Richardson and Roberts [34]. The breakpoint (BP) value was defined as the last ratio attained by the rat over the 60 min session. GSK4716 was administered IP 30 min before the start of the PR test at the doses of 0, 2.5, and 5 mg/kg.

#### Spontaneous locomotor activity

Acute effect of GSK4716 administration was assessed on spontaneous locomotor activity in male alcohol-naive Wistar rats. Rats were randomly injected with either 0, 1.25, 2.5 and 5 mg/kg GSK4716 30 min before the start of the test and then placed into an arena provided with infrared photo beams (Med Associates, USA). The rat locomotor activity was measured over a 30-min session.

## Results

### Systems biology identification of candidate drivers of alcohol dependence-associated gene network dysregulations in the human PFC

The *corto* algorithm and pipeline [4] was used to perform MRA using a human dataset comprising PFC gene expression profiles from 68 individuals with histories of alcohol dependence vs. 60 matched control samples. A PFC gene regulatory network was reconstructed from the human GTEX Consortium frontal cortex dataset of 129 human samples [26] using *corto* and a collection of candidate master regulators (MRs) consisting of a curated list of human transcription factors (TFs) derived from Gene Ontology category DNA-binding transcription factor activity (GO:0003700). Fig. 1A shows the top 10 ranking MRs identified as drivers of PFC gene signatures associated with a history of alcohol dependence including ESRRG (ERRγ). The normalized enrichment score (NES) of each candidate MR, as well as the p-value and the Benjamini-Hochberg corrected p-value are shown in Supplementary table 1. To better define the gene regulatory network in which ESRRG is active we performed a correlation analysis between genetical and transcriptomical events using a modified DIGGIT pipeline [27]. Genes affecting the activity of ESRRG in at least 3 or 2 datasets with a corrected p-value threshold of 0.01 are shown in Fig 1B and 1C, respectively.

**Fig 1.**
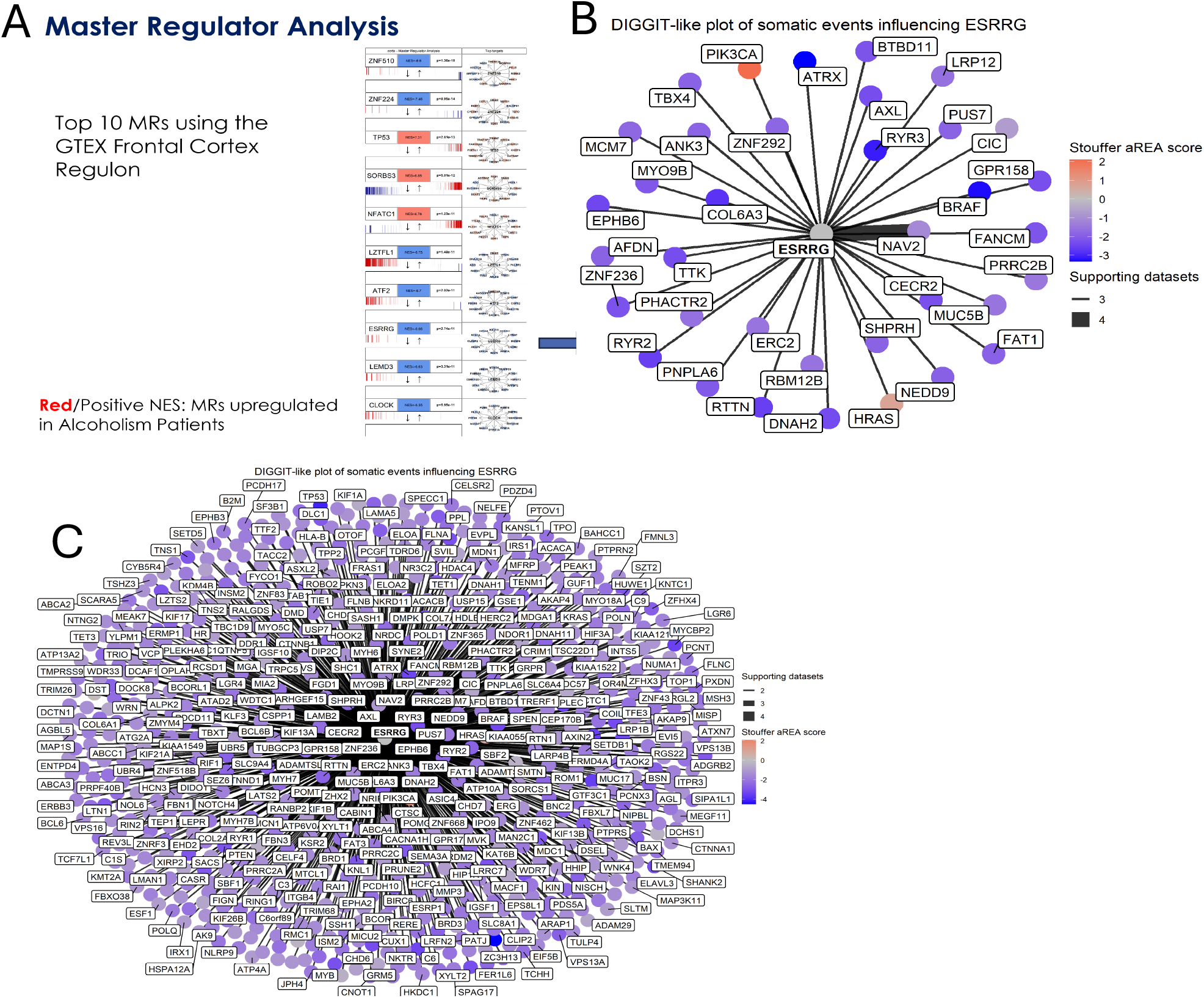
ESRRG (ERRγ) is a master regulator gene (MR) governing the gene network dysregulations in the pre-frontal cortex (PFC) of individuals with a history of alcohol dependence. **A)** High-ranking MRs identified by master regulator analysis (MRA) of PFC gene expression profiles from 68 individuals with histories of alcohol dependence vs. 60 matched control samples. **B-C)** We detected 36 genes whose somatic mutations are statistically significantly correlated with ESRRG activity in at least 3 datasets, p<0.001 (B); mutations in 711 genes were correlated with ESRRG activity in at least 2 datasets (C); and 3,665 genes in at least one dataset (not shown).

### Mice Experiments

#### Effects of the GSK4716 on binge-like alcohol drinking in the DID paradigm in C57BL/6J mice

Acute administration of GSK4716 resulted in a statistically significant reduction of alcohol intake (1-way ANOVA: F(3,40)= 7.29; p<0.0005) in the mouse groups treated with 2.5 and 5 mg/kg in comparison to the mouse vehicle-treated group (**p<0.005; ****p<0.0001 by Tukey’s post hoc test) during the first 2 h of the drinking session (Fig 2A). The alcohol-reducing effect of GSK4716 persisted through the entire 4 h session at 2.5 and 5 mg/kg GSK4716 doses [1-way ANOVA: F (3,40) = 5.86; p<0.005; **p<0.005 by Tukey’s post hoc test] (Fig 2B).

**Fig 2.**
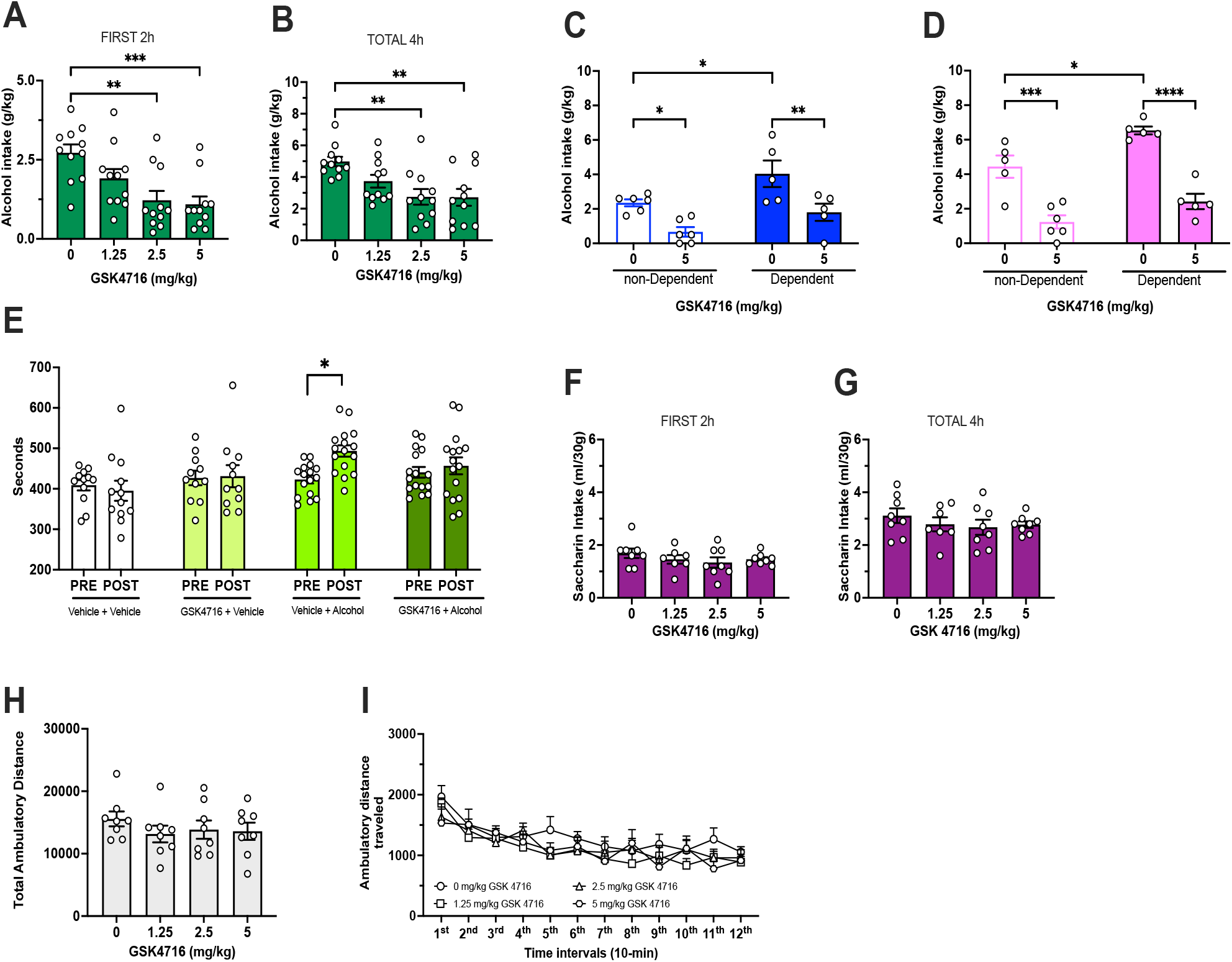
GSK4716 selectively reduced alcohol consumption and alcohol reward properties without affecting saccharin intake or spontaneous locomotor activity in C57BL/6J mice. GSK4716 (2.5 and 5 mg/kg) significantly reduced binge-like alcohol consumption during the first 2 h **(A)** and over the entire 4 h session **(B)** (**p < 0.005, ****p < 0.0001 by Tukey’s test. GSK4716 (5 mg/kg) significantly suppressed alcohol consumption in both male **(C)** and female mice **(D)** (*p < 0.05, **p < 0.01, ***p < 0.001, ****p < 0.0001, by Šidák test). **E)** GSK4716 (5 mg/kg) reduced alcohol reward properties (*p< 0.05, by Šidák test). GSK4716 did not affect saccharin consumption during the first 2 h **(F)** or over the 4 h session **(G)**. GSK4716 did not affect spontaneous locomotor activity, as measured by **(H)** total ambulatory distance over 2 h or **(I)** distance across twelve 10-min intervals.

#### Effect of GSK47162 on 2-bottle choice after CIE

##### Males

Two-way repeated measure (RM) ANOVA of alcohol intake (g/kg) during the limited 2BC session revealed a significant main effect of alcohol vapor exposure (F (1,18) = 9.59; p<0.01) and treatment (F (1,18) = 18.52; p<0.0005). Post hoc analysis revealed that dependent mice consumed more alcohol than non-dependent mice (4.04 g/kg vs 2.35 g/kg) (*p<0.05 by Šidák post hoc test). Administration of GSK4716 (5 mg/kg) drastically reduced alcohol intake in both groups (0.67 g/kg vs 1.8 g/kg, respectively) (*p< 0.05, **p<0.01 Šidák post hoc test) (Fig 2C).

##### Females

Two-way RM ANOVA of alcohol intake (g/kg) during the limited 2BC session revealed a significant main effect of alcohol vapor exposure (F(1,17) = 13.48; p<0.005) and treatment (F(1, 1) = 67.11; p<0.0001). Post hoc analysis showed that dependent mice consumed more alcohol than dependent (6.53 g/kg vs. 4.44 g/kg, respectively) (*p<0.05 by Šidák post hoc test). Administration of GSK4716 significantly reduced alcohol consumption in both non-dependent (***p<0.05) and dependent (****p<0.0001) mouse groups compared to their vehicle-treated control groups (Šidák post hoc test) (Fig 2D).

#### Effect of GSK4716 on conditioned place preference

Three-way ANOVA indicated a significant main effect of treatment (F (1,52) = 7.83, p<0.01)) but not main effect of conditioning (F (1,52) = 2.008, p>0.05)) or pretreatment (F (1,50) = 0.58, p<0.01)). Post hoc analysis indicated a significant pre-post conditioning difference in the vehicle + alcohol-treated group (*p< 0.05 by Šidák post hoc test) (Fig. 2E).

#### Effect of GSK4716 on saccharine drinking in C57BL/6J mice

No significant effect of GSK4716 administration on saccharine intake (ml/30g) was observed at any dose during the first 2 h (1-way ANOVA: F(3,27) = 0.86; p>0.05) (Fig 2F) or over the entire 4 h session (1-way ANOVA: F(3,27) = 0.61; p>0.05) (Fig 2G).

#### Effect of GSK4716 on mouse spontaneous locomotor activity

One-way ANOVA revealed no significant differences among groups treated with either GSK4716 or vehicle on total ambulatory distance over a 2 h session (F(3,28) = 0.61; p>0.05) (Fig 2H). In addition, 2-way RM ANOVA of ambulatory distance across twelve 10-min intervals, revealed a significant effect of time (F(11,308) = 15.35; p<0.0001) (Fig. 2I).

### Rat experiments

#### Effect of GSK4716 on FR1 schedule of reinforcement

##### Number of responses on the alcohol lever

Alcohol dependent rats exhibited significantly higher responding on the alcohol lever compared to non-dependent rats (31.7 *vs* 55.6 lever presses over 30 min session; main effect of group: F(1,52) =18.20; p<0.0001). Two-way ANOVA revealed that GSK4716 administration significantly reduced alcohol self-administration in both non-dependent and dependent rat groups (main effect of treatment: F(2,52) =8.23; p<0.005). Post-hoc analysis indicated that only the 5 mg/kg dose of GSK4716 significantly reduced alcohol lever-responses in both non-dependent and dependent rat groups (**p<0.005; *p<0.05 by Tukey’s post hoc test) (Fig 3A).

**Fig 3.**
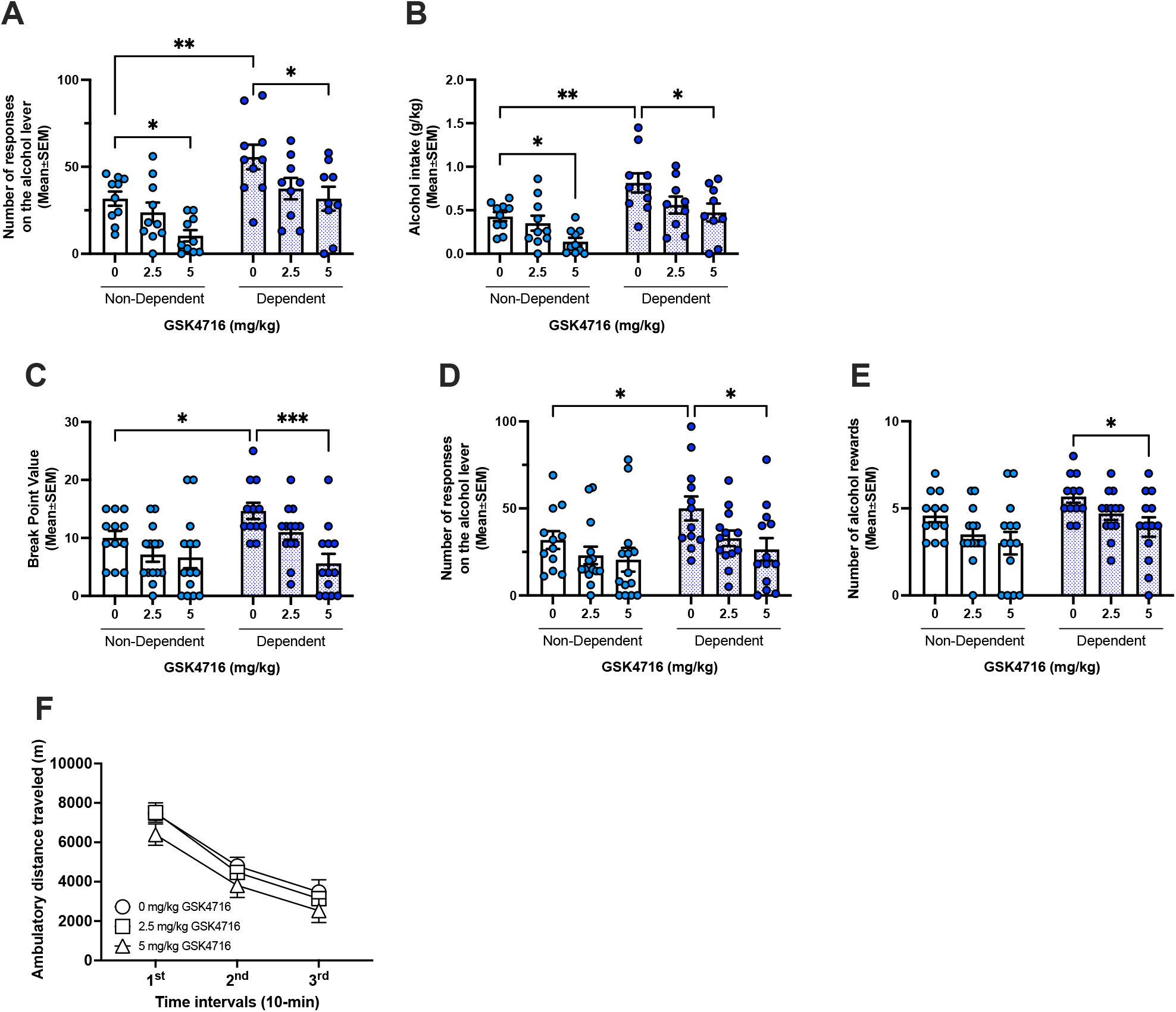
GSK4716 (5 mg/kg) reduced alcohol self-administration without affecting spontaneous locomotor activity in Wistar rats. Alcohol-dependent rats exhibited higher responding on the alcohol lever and greater alcohol intake compared with non-dependent rats. GSK4716 significantly suppressed **(A)** number of responses on the alcohol lever and **(B)** alcohol intake (*p < 0.05, **p < 0.005, by Tukey’s test). Similarly, GSK4716 reduced break point **(C)**, number of responses on the alcohol lever **(D)** and number of alcohol rewards **(E)** (*p < 0.05, ***p < 0.001, by Tukey’s test). **F)** GSK4716 did not affect spontaneous locomotor activity of alcohol-naive rats.

##### Amount of self-administered alcohol

Similarly, alcohol dependent rats self-administered greater amounts of alcohol than non-dependent rats (0.43 *vs* 0.81 g/kg/30 min session; main effect of group: F(1, 52) = 19.90; p<0.0001). Two-way ANOVA revealed a significant main effect of GSK4716 treatment on alcohol intake (F (2,52) = 6.84; p<0.005). Post-hoc comparisons revealed that only the 5 mg/kg dose of GSK4716 significantly decreased alcohol self-administration in both non-dependent and dependent rat groups (**p<0.005; *p<0.05, by Tukey’s post hoc test) (Fig 3B).

#### Effect of GSK4716 on a PR schedule of reinforcement

##### Break Point (BP) Value

Alcohol dependent rats exhibited a significantly higher motivation to obtain alcohol compared to non-dependent rats, as indicated by higher BP values (10 *vs* 14.67). Two-way ANOVA revealed significant main effects of both groups (F(1,72) = 4.25; p<0.05)) and treatment (F(2,72) = 8.57; p<0.001). Post-hoc analysis showed that only the 5 mg/kg dose of GSK4716 significantly reduced the motivation for alcohol only in the dependent rat group (***p <0.0005; *p<0.05 by Tukey’s post hoc test) (Fig 3C).

##### Number of responses on the active lever

Responding on the active lever was significantly higher in the alcohol dependent rat group than the non-dependent rat group (31.8 vs 50 responses over 60-min session). Two-way ANOVA revealed significant main effects of group (F(1,72) = 5.49; p<0.05) and treatment (F(2,72) = 4.53; p<0.05). Post-hoc analysis revealed that the 5 mg/kg dose of GSK4716 significantly reduced lever-responses for alcohol in the dependent rat group only (*p<0.05 by Tukey’s post hoc test) (Fig 3D). Lastly, GSK4716 reduced the number of alcohol rewards. Two-way ANOVA revealed significant main effects of group (F(1,72) = 7.47; p<0.001) and treatment (F(2,72) = 46.01; p<0.05). Post-hoc analysis indicated that 5 mg/kg of GSK4716 significantly reduced the number of alcohol rewards in the dependent rat group (*p<0.05 by Tukey’s post hoc test) with a trend toward reduction in the non-dependent rat group (p=0.056) (Fig 3D).

#### Effect of GSK4716 on spontaneous locomotor activity

One-way ANOVA analysis of the total ambulatory distance traveled by rats during a 30-minute session showed that administration of GSK4716 did not significantly affect locomotor activity (F(2,27) = 0.89; p>0.05). Furthermore, two-way RM ANOVA of ambulatory distance across three 10-min intervals revealed a significant effect of time (F(2,81) = 52.63; p<0.0001) (Fig. 3G).

## Discussion

The pathogenesis of AUD is complex, involving neurobiological, genetic, and environmental factors [35]. These factors interact to cause long-lasting changes in brain circuits related to reward, motivation, and self-control, leading to compulsive alcohol use despite negative consequences [35]. Here, we used a systems biology approach to reverse engineer the transcriptional regulation network from the PFC of individuals with histories of alcohol dependence to identified unbiased candidate Master Regulators (MRs) that contribute to their phenotypic behavioral manifestations.

We identified the estrogen-related receptor gamma (ERRγ) gene ESRRG as a high-ranking MR and we selected it for functional validation due to its translational and druggability potential [24, 25].

In the present study, we demonstrate that acute administration of the ERRγ agonist GSK4716 reduces alcohol consumption in mice in the drinking in the dark (DID) paradigm of binge drinking, as well as in both dependent (via chronic intermittent vapor exposure, CIE) and non-dependent mice. Similarly, GSK4716 reduced the reinforcing and motivational properties of alcohol in both non-dependent and dependent rats under fixed and progressive ratio schedules of reinforcement. Importantly, GSK4716 did not affect saccharin intake or spontaneous locomotor activity, indicating that its effects are specific to alcohol-related behaviors rather than a general suppression of reward or motor activity. Moreover, GSK4716 prevented the development of alcohol-induced conditioned place preference in alcohol-naive mice, further supporting the involvement of ERRγ in the reward-brain circuit.

The central nervous system (CNS) is one of the most energy-demand organs in the body, and mitochondria, often referred as “the powerhouse of the cells,” play a crucial role in sustaining its function [36, 37]. Accordingly, mitochondrial dysfunction has been implicated in the onset and progression of several brain diseases, including neurodegenerative conditions such as Alzheimer’s, Parkison’s, and Huntington’s disease as well as various psychiatric diseases [36]. A growing body of literature demonstrates that excessive and chronic alcohol consumption impair mitochondria function, which in turn may contribute to neuroinflammation and the development of AUD [38]. For instance, binge drinking during adolescence disrupts mitochondrial bioenergetic processes with deficits persisting into adulthood [39]. Preclinical studies have shown that mitochondria in the PFC of C57BL/6J mice exposed to four cycles of chronic intermittent exposure to ethanol vapors (CIE) exhibit altered morphology, reduced respiratory capacity, increased expression of fission proteins, and decreased levels of fusion protein and morphological changes akin to AD [40].

ERRγ is a nuclear receptor known to be involved in metabolism and metabolic disease [41]. Loss of ERRγ impairs neuronal metabolic capacity and long-term potentiation deficits observed in ERRγ^−/−^ hippocampal slices, which can be rescued by the mitochondrial oxidative phosphorylation substrate pyruvate [42]. In line with these findings, mice lacking neuronal ERRγ in the cerebral cortex and hippocampus exhibit defects in spatial learning and memory as demonstrated by poorer performance in the Morris water maze test compared to control mice [42]. Thus, we hypothesize that activating ERRγ by administration of its agonist GSK4716 may improve mitochondrial activity, particularly in the brain reward circuits, therefore limiting alcohol taking and seeking behavior.

In conclusion, we identified the ERRγ gene (ESRRG) as a high-ranking master regulator contributing to the phenotypic behavior of individuals with histories of alcohol dependence. We further validated its role in multiple established rodent models of excessive alcohol consumption and provided the first evidence that pharmacological activation of ERRγ interferes with the reinforcing and motivational properties of alcohol, supporting ERRγ as a promising therapeutic target for AUD.

## Funding

This work was supported by NIH Grant AA021667; IL was partially supported by training grant # T32 AA007456.

**All authors approved the final version of the manuscript**

